# Prediction of protein mutational free energy: benchmark and sampling improvements increase classification accuracy

**DOI:** 10.1101/2020.03.18.989657

**Authors:** Brandon Frenz, Steven Lewis, Indigo King, Hahnbeom Park, Frank DiMaio, Yifan Song

**Affiliations:** Cyrus Biotechnology; University of Washington

## Abstract

Software to predict the change in protein stability upon point mutation is a valuable tool for a number of biotechnological and scientific problems. To facilitate the development of such software and provide easy access to the available experimental data, the ProTherm database was created. Biases in the methods and types of information collected has led to disparity in the types of mutations for which experimental data is available. For example, mutations to alanine are hugely overrepresented whereas those involving charged residues, especially from one charged residue to another, are underrepresented. ProTherm subsets created as benchmark sets that do not account for this often underrepresented certain mutational types. This issue introduces systematic biases into previously published protocols’ ability to accurately predict the change in folding energy on these classes of mutations. To resolve this issue, we have generated a new benchmark set with these problems corrected. We have then used the benchmark set to test a number of improvements to the point mutation energetics tools in the Rosetta software suite.

## Introduction

The ability to accurately predict the stability of a protein upon mutation is important for numerous problems in protein engineering and medicine including stabilization and activity optimization of biologic drugs. To perform this task a number of strategies and force fields have been developed, including those that perform exclusively on sequence(Capriotti et al.; Casadio et al.) (Kumar et al.) as well as those that involve sophisticated physical force fields both knowledge based(Sippl) (Gilis and Rooman) (Potapov et al.), physical models (Pitera and Kollman) (Benedix et al.) (Pokala and Handel), and hybrids (Guerois et al.) (Pitera and Kollman; Park et al.) (Kellogg et al.) (Quan et al.) (Jia et al.).

To facilitate the development of these methodologies and provide easy access to the available experimental information the ProTherm database(Uedaira et al.) was developed. This database collects thermodynamic information on a large number of protein mutations and makes it available in an easy to access format. At the time of this writing it contains 26,045 entries.

Due to its ease of access the ProTherm has served as the starting point for a number of benchmark sets used to validate different stability prediction software packages, including those in the Rosetta software suite. However significant biases exist in the representation of different types and classes of mutations in the ProTherm, as it is derived from the existing literature across many types of proteins and mutations. The most obvious example of this is the large number of entries involving a mutation from a native residue to alanine as making this type of mutation is a common technique used to find residues important for protein function. Therefore a large number of the benchmark sets derived from the ProTherm, which did not account for this bias, have significantly under or overrepresented these classes of mutations. These findings suggest previous reports on the accuracy of stability prediction software does not accurately reflect these tools’ ability to predict stability changes across all classes of mutations.

To address this issue we have generated a novel benchmark subset which accounts for this bias in the database (Supplemental Table 1). We then used this benchmark set to validate and improve upon an existing free energy of mutation tool within the Rosetta software suite, “Cartesian ddG,” first described in Park et al. 2016. (Park et al.). We have also brought in new algorithms originally developed for cryoEM refinement to help improve sampling around difficult mutations involving proline. (Wang et al.)

## Results

In order to benchmark our Rosetta-based stability prediction tools we classified the possible mutations into 17 individual categories as well as reported results on three aggregate categories. We analyzed five previously published benchmark sets to determine their coverage across the different classes of mutations and found them inadequate in a number of categories, especially involving charged residues (Figure 1 A-E). For example, the number of data points for mutational types ranged from 0-24 for negative to positive, 0-50 for positive to negative, 3-28 for hydrophobic to negative, and 3-44 for hydrophobic to positive entries across the benchmark sets tested. Mutations to and/or from hydrophobic residues dominated the benchmark sets ranging from 75%-92% of the total entries.

**Figure 1.**
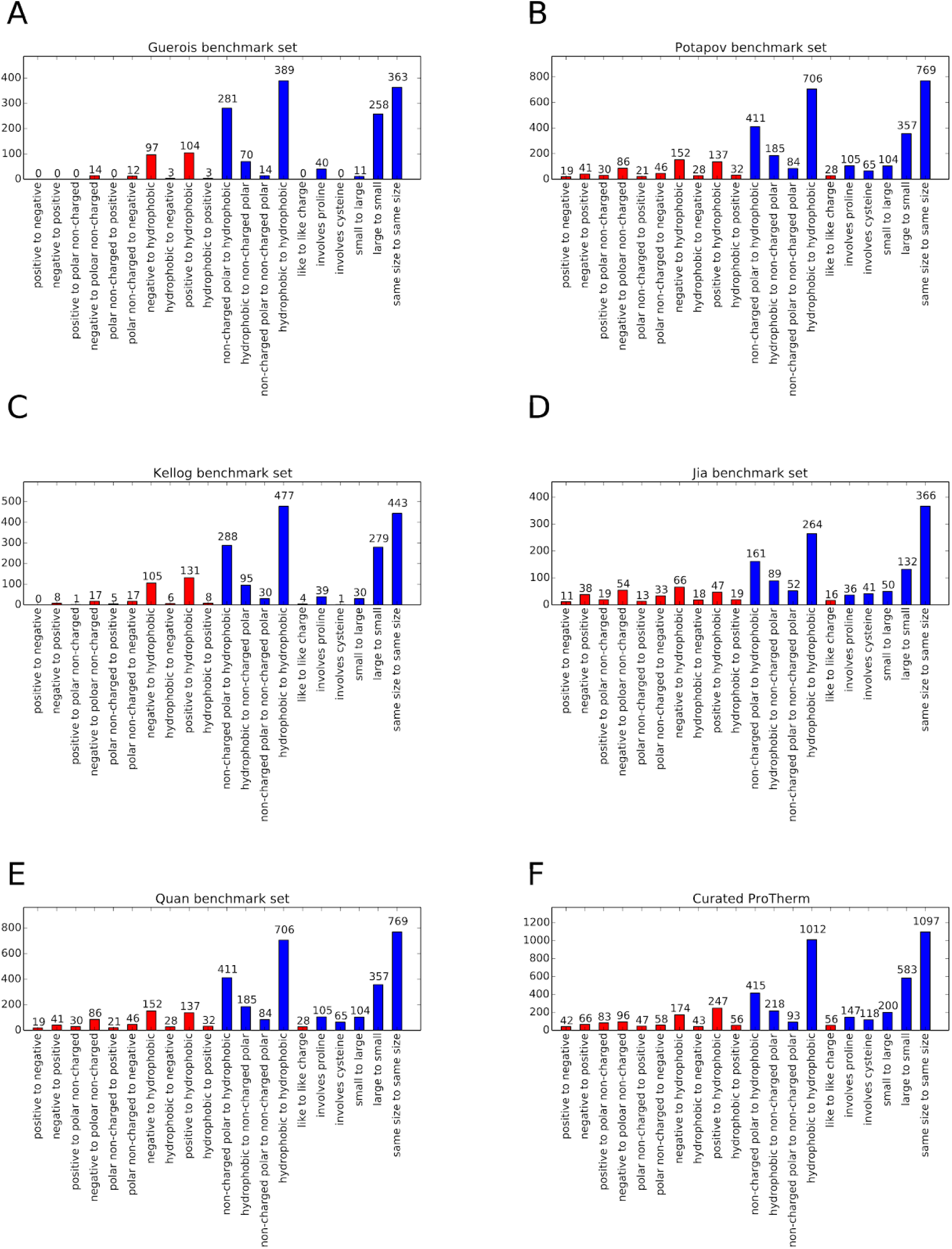
Benchmark Set mutational category statistics. This figure shows the population of different mutation classes used to benchmark a number of methods predict the change in free energy upon mutation the citations for these benchmark sets are as follows A. Guerois (Guerois et al.) B. Potapov ((Potapov et al.) C. Kellog (Kellogg et al.) D. Jia (Jia et al.) E. Quan (Quan et al.). As well as the overall representation of the curated ProTherm (F). Classes involving charged residues are colored in red. All data sets are significantly biased in their types of mutations present.

To compare the composition of these benchmark sets to that of the database we examined the curated ProTherm (ProTherm*)) provided by Kortemme et al. (https://guybrush.ucsf.edu/benchmarks/benchmarks/DDG) which is a selection of entries containing only mutations which occur on a single chain and provide experimental ΔΔG values. We find that significant biases still exist here, with several categories having fewer than 50 unique mutations. These include: positive to negative, 42; hydrophobic to negative, 43; and non-charged polar to positive, 47. Mutations involving hydrophobic residues are still overrepresented with 74% of all mutations in the database involving hydrophobic residues (Figure 1 F).

To sample more broadly across all types of mutations and remove sources of bias in our algorithm training we created a new benchmark set of single mutations that are more balanced across mutational types and avoid other biases. To generate this set we performed the following operations:

1. Removed any entries from the curated ProTherm* that occur on the interface of a protein complex or interact with ligands -- the energetics of these mutations would include intra-protein and inter-molecular interactions that would alter the desired intra-protein energetics of a free energy calculation.
2. We removed entries of identical mutation on similar-sequence (> 60%) backbones. For mutations occurring at the same position in similar sequences, if the mutation is identical (e.g. L->I) and the sequence identity > 60%, then that mutation is included only once in the database; if the mutation is not identical (L-->I in one protein and L-->Q in another) then the mutation is included.
3. We populated each mutually exclusive mutation category with 50 entries except for the cases where insufficient experimental data points exist.
4. When multiple experimental values (including identical mutations as identified in point 2 above) were available we chose the ΔΔG value taken at the pH closest to 7.

The resulting benchmark set contains 767 entries across a range of different types and classes of mutations (Figure 2).

**Figure 2.**
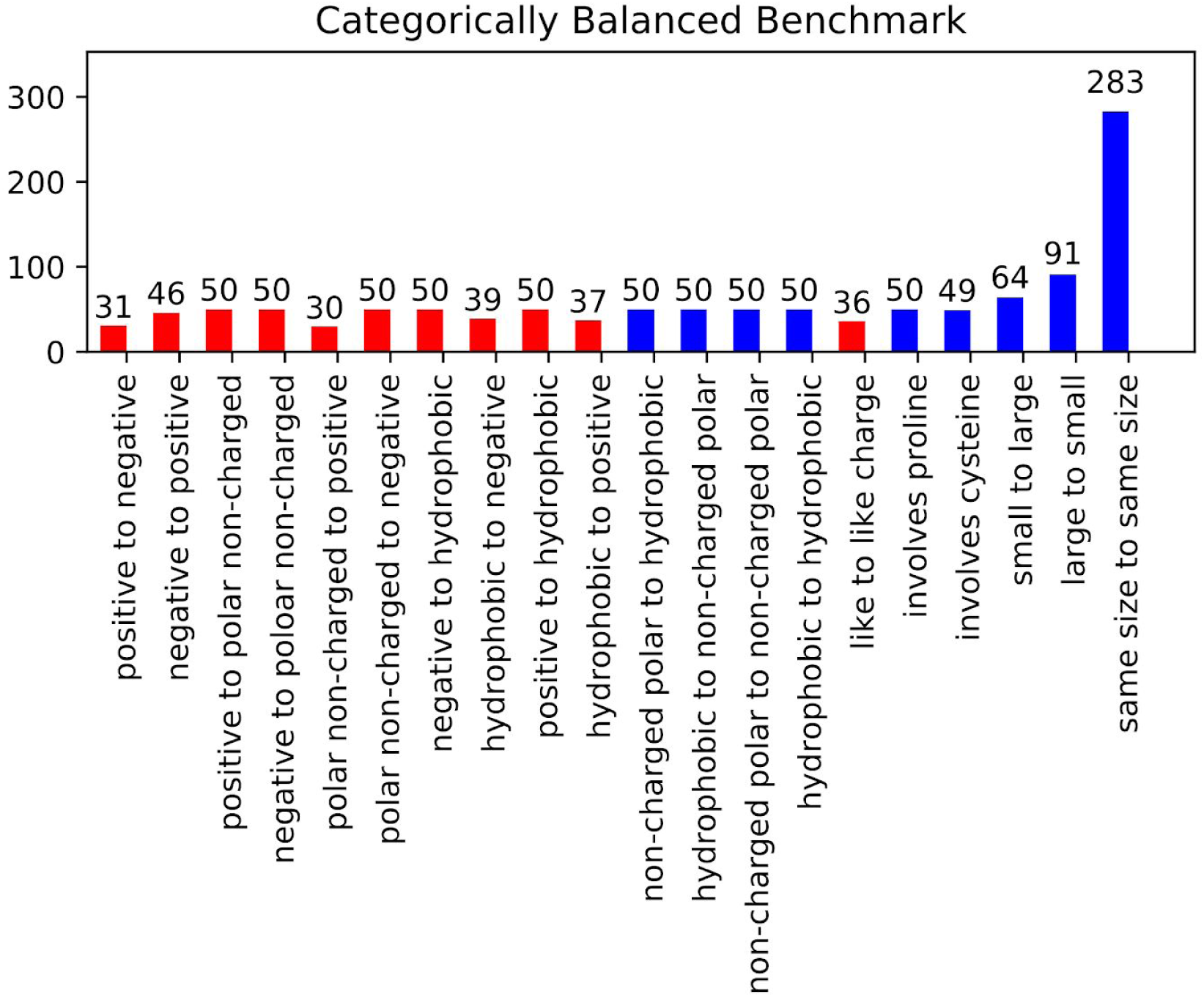
Categorically Balanced Benchmark mutational category statistics. This figure shows the metrics of our new benchmark set selected to provide a more balanced representation of different mutation classifications. Classes involving charged residues are shown in red.

We tested Protocol 3 described in Kellog et al. (Kellogg et al.) on this benchmark set and found that prediction ability varied widely across different mutational classes (Table 1). This suggests that reported correlation metrics of different protocols may be heavily influenced by the composition of their benchmark sets. It also suggests that certain classes of mutations may pose a more difficult prediction problem.

**Table 1.** Correlations and Predictive Index for Protocol 3 and our improved Cartesian ddG across different mutation categories. This table contains the Pearson’s R correlations for each class of mutations in our benchmark set for both Protocol 3 and Cartesian ddG. Each is repeated three times using the same inputs and the average and standard deviation are shown. Given the sensitivity to outliers of Pearson’s R we also report it as Pearson’s R filtered after removing up to 5 outliers from each set. An outlier is defined as any single entry which when removed changes the correlation coefficient by 0.025 or greater. We also report the predictive index which is less sensitive to the absolute free energy of a prediction but rather whether it can be correctly classified. Cartesian ddG significantly outperforms protocol 3 both in the unfiltered Pearson’s R and in Predictive Index.

|  | Protocol 3 |  |  | Cartesian DDG |  |  |
| --- | --- | --- | --- | --- | --- | --- |
| Mutation Type | Pearson's R | Pearson's R Filtered | Predictive Index | Pearson's R | Pearson's R Filtered | Predictive Index |
| small to large | 0.54±0.000 | 0.68±0.000 | 0.57±0.001 | 0.53±0.060 | 0.67±0.073 | 0.59±0.053 |
| large to small | 0.56±0.018 | 0.76±0.000 | 0.59±0.000 | 0.60±0.035 | 0.81±0.029 | 0.70±0.021 |
| positive to negative | 0.40±0.000 | 0.79±0.000 | 0.28±0.000 | 0.58±0.014 | 0.81±0.044 | 0.69±0.026 |
| negative to positive | 0.34±0.000 | 0.61±0.000 | 0.26±0.000 | 0.35±0.014 | 0.54±0.037 | 0.42±0.032 |
| negative to hydrophobic | 0.27±0.000 | 0.55±0.000 | 0.27±0.000 | 0.56±0.038 | 0.66±0.042 | 0.60±0.0361 |
| hydrophobic to negative | 0.83±0.000 | 0.87±0.000 | 0.84±0.000 | 0.77±0.021 | 0.83±0.025 | 0.84±0.018 |
| positive to hydrophobic | -0.06±0.000 | 0.23±0.000 | 0.01±0.000 | 0.51±0.060 | 0.62±0.040 | 0.55±0.052 |
| hydrophobic to positive | 0.57±0.000 | 0.73±0.000 | 0.63±0.002 | 0.54±0.040 | 0.74±0.044 | 0.73±0.023 |
| non-charged polar to positive | 0.40±0.000 | 0.67±0.000 | 0.39±0.004 | 0.27±0.136 | 0.69±0.052 | 0.35±0.128 |
| positive to non-charged polar | 0.32±0.000 | 0.67±0.000 | 0.52±0.000 | 0.46±0.019 | 0.77±0.036 | 0.67±0.012 |
| non-charged polar to negative | 0.64±0.000 | 0.73±0.000 | 0.69±0.000 | 0.49±0.075 | 0.72±0.053 | 0.52±0.092 |
| negative to non-charged polar | 0.13±0.000 | 0.44±0.000 | -0.07±0.000 | 0.39±0.059 | 0.66±0.014 | 0.43±0.051 |
| non-charged polar to hydrophobic | 0.70±0.000 | 0.70±0.000 | 0.64±0.001 | 0.64±0.048 | 0.78±0.006 | 0.57±0.038 |
| hydrophobic to non-charged polar | 0.41±0.000 | 0.66±0.000 | 0.39±0.000 | 0.61±0.065 | 0.76±0.017 | 0.63±0.089 |
| non-charged polar to non-charged polar | 0.76±0.000 | 0.76±0.000 | 0.66±0.002 | 0.66±0.029 | 0.78±0.013 | 0.74±0.028 |
| hydrophobic to hydrophobic | 0.67±0.000 | 0.74±0.000 | 0.72±0.000 | 0.42±0.033 | 0.70±0.037 | 0.62±0.025 |
| charge to charge | 0.31±0.000 | 0.73±0.000 | 0.36±0.000 | 0.22±0.057 | 0.67±0.064 | 0.29±0.072 |
| involves cysteine | 0.25±0.000 | 0.63±0.000 | 0.27±0.000 | 0.05±0.015 | 0.42±0.040 | 0.11±0.034 |
| involves proline | 0.02±0.000 | 0.54±0.000 | 0.33±0.000 | 0.44±0.048 | 0.73±0.024 | 0.50±0.033 |
| same size | 0.36±0.000 | 0.38±0.000 | 0.37±0.000 | 0.36±0.013 | 0.46±0.011 | 0.46±0.007 |
| everything | 0.25±0.000 | 0.47±0.000 | 0.48±0.000 | 0.45±0.009 | 0.45±0.009 | 0.58±0.004 |

To address some of these poor performing metrics a number of changes were made to the more modern Rosetta ΔΔG protocol, Cartesian ddG, from Park et al. 2016 (Park et al.). These include changes to the preparation step of the input model and increased optimization of the backbone around residues that are being mutated to or from proline (Figure 3). To improve the preparation step (step 1), we tested Cartesian Relax (as opposed to traditional torsion space Relax) both with and without all atom constraints and found that correlations were better without constraints (data not shown). This likely has to do with the use of Cartesian minimization during step 4, and the importance of preparing a structure with similar sampling methods to those used during mutational energy evaluation. In considering the final energy for a mutation, we also compared taking the average of three experiments vs. a multi-run convergence criterion and settled on the convergence criterion method. We also increased backbone sampling for mutations involving proline taking advantage of new methods developed for CryoEM refinement. (Wang et al.) We have also refactored the Cartesian ddG code for efficiency and modifiability.

**Figure 3.**
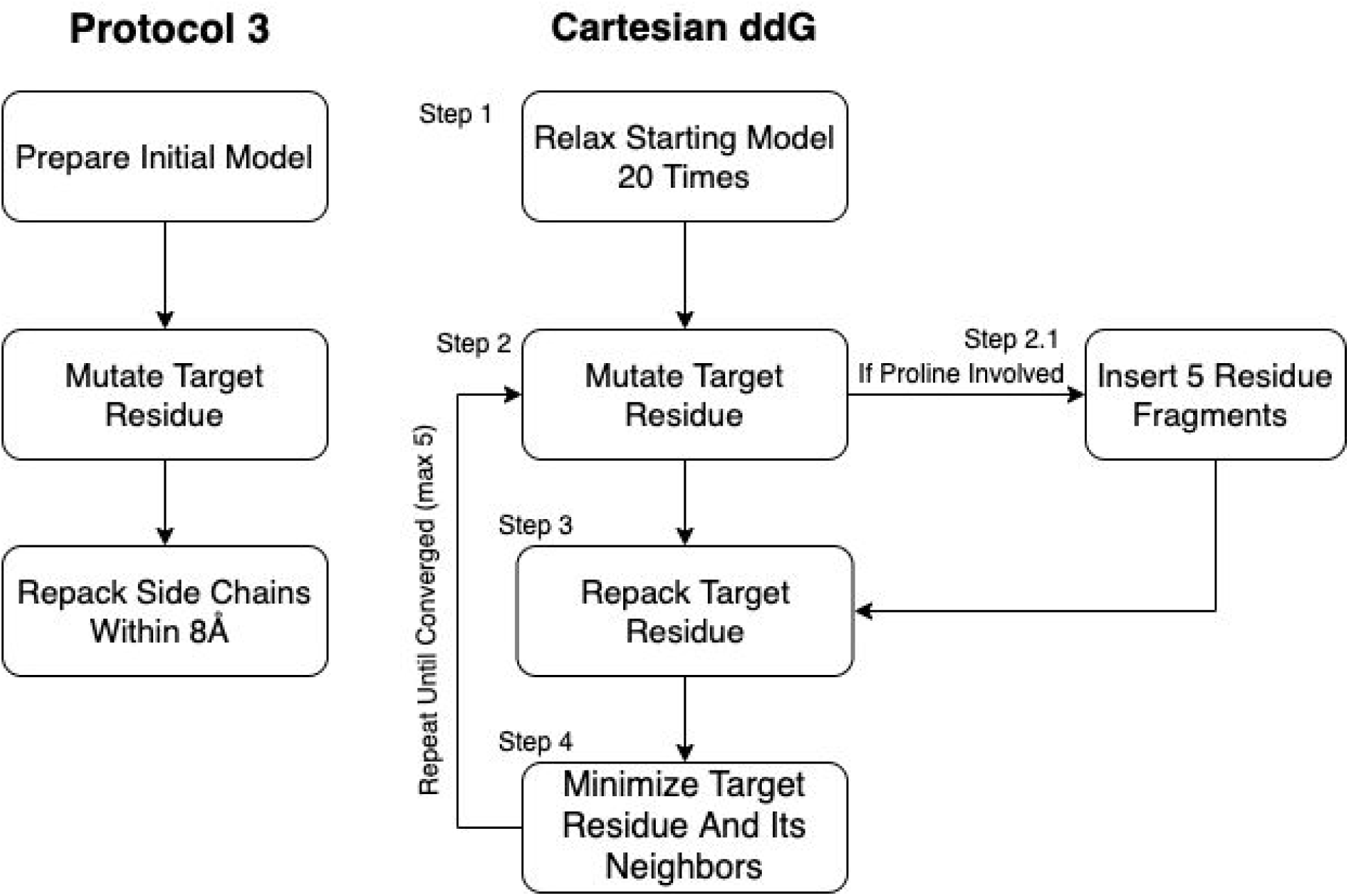
Diagram of “Protocol 3” and Cartesian ddG. This figure diagrams the steps involved in the older protocol 3 as well as the Cartesian ddG protocol. Novel changes described in this paper include the removal of constraints during step 1 of the cartesian ddG protocol, the addition of Step 2.1 for mutations involving proline as well as the choice to repeat testing until the protocol converges on a lower energy score instead of a fixed number (3) of times.

In addition to Pearson’s R we analyzed prediction power by classification errors instead of by correlation. A mutation is classified as stabilizing if the change in free energy is ≤-1 kcal/mol, it is classified as destabilizing if the change is ≥1 kcal/mol, and neutral if it falls between these values. Each mutation is assigned a value of 0 for destabilizing, 1 for neutral, and 2 for stabilizing. We then scored each entry by taking the absolute value of the difference between the assigned value for the experiment and the prediction. A value of 0 indicates the prediction was correct, 1 indicates the prediction was moderately incorrect, i.e. the mutation is destabilizing and the prediction was neutral, and 2 indicates the prediction was egregiously wrong.

This new and optimized Cartesian ddG generally improves performance overall especially in the ability to accurately classify mutations as shown by the Predictive Index (Pearlman and Charifson) and by the large reduction egregious errors in classification (Tables 1, 2). For example, the number of mutations predicted as stabilizing when they are destabilizing or vice versa fell from an average of 53 with Protocol 3 to an average of 31.3 across 3 replicates. “Off by 1” errors are also lower (317.3 vs 292.7). This trend is much stronger than the improvement in correlations, and more importantly reflects the practical value of correctly classifying mutational categories. For example in protein engineering, a protein designer’s practical interest is whether any given mutation is stabilizing at all, more than which of two mutations is more stabilizing.

**Table 2.** The ability of Protocol 3 and Cartesian ddG to correctly classify mutations. This table shows the ability of Protocol 3 and Cartesian ddG to correctly classify a mutation. Mutations are assigned a value of 0 for destabilizing, 1 for neutral, and 2 for stabilizing. The absolute value of the difference between the predicted class and the experimental class summed across the benchmark and the counts are reported here. The average of 3 replicates and their standard deviations are shown. Cartesian ddG correctly classifies far more entries 441 vs 396, and produces fewer mis classifications, especially “Off By Two” errors, 31.3 v 53.

| mutation class | Protocol 3 |  |  | Cartesian ddG |  |  |
| --- | --- | --- | --- | --- | --- | --- |
|  | Same Class | Off By One | Off By Two | Same Class | Off By One | Off By Two |
| small to large | 35.7±0.6 | 27.3±0.6 | 1±0 | 39.7±4.6 | 21.7±4.0 | 2.7±0.6 |
| large to small | 55.0±0.0 | 31.0±0.0 | 5±0 | 50.3±2.5 | 37.3±2.5 | 3.0±0.0 |
| positive to negative | 10.0±0.0 | 19.0±0.0 | 2±0 | 21.3±1.2 | 9.7±1.2 | 0.0±0.0 |
| negative to positive | 12.0±0.0 | 26.0±0.0 | 8±0 | 25.7±1.2 | 17.0±1.7 | 3.3±0.6 |
| negative to hydrophobic | 16.0±0.0 | 25.0±0.0 | 9±0 | 27.7±2.1 | 17.7±2.1 | 4.7±1.2 |
| hydrophobic to negative | 21.0±0.0 | 17.0±0.0 | 1±0 | 25.0±1.0 | 13.7±1.2 | 0.0±0.0 |
| positive to hydrophobic | 25.0±0.0 | 21.0±0.0 | 4±0 | 25.0±3.5 | 23.7±2.1 | 1.0±1.0 |
| hydrophobic to positive | 27.0±0.0 | 7.0±0.0 | 3±0 | 28.7±1.5 | 7.3±1.5 | 1.0±0.0 |
| non-charged polar to positive | 15.0±0.0 | 15.0±0.0 | 0±0 | 14.3±1.2 | 14.0±1.0 | 1.7±0.6 |
| positive to non-charged polar | 29.0±0.0 | 19.0±0.0 | 2±0 | 33.0±3.6 | 15.3±4.0 | 1.3±0.6 |
| non-charged polar to negative | 24.0±0.0 | 24.0±0.0 | 2±0 | 27.0±2.6 | 20.3±1.5 | 2.7±1.2 |
| negative to non-charged polar | 22.0±0.0 | 18.0±0.0 | 10±0 | 24.3±2.1 | 22.7±1.2 | 2.3±0.6 |
| non-charged polar to hydrophobic | 24.0±0.0 | 24.0±0.0 | 2±0 | 23.0±4.0 | 25.3±4.0 | 1.7±0.6 |
| hydrophobic to non-charged polar | 28.0±0.0 | 21.0±0.0 | 1±0 | 32.0±1.0 | 16.0±0.0 | 2.0±1.0 |
| non-charged polar to non-charged polar | 28.7±0.6 | 19.3±0.6 | 2±0 | 32.0±1.0 | 15.7±0.6 | 2.3±0.6 |
| hydrophobic to hydrophobic | 38.0±0.0 | 11.0±0.0 | 1±0 | 36.3±0.6 | 13.0±1.0 | 0.7±0.6 |
| charge to charge | 23.0±0.0 | 12.0±0.0 | 0±0 | 14.7±1.2 | 20.0±1.7 | 0.3±0.6 |
| involves cysteine | 27.0±0.0 | 20.0±0.0 | 2±0 | 21.7±0.6 | 23.0±1.7 | 4.3±2.1 |
| involves proline | 27.0±0.0 | 19.0±0.0 | 4±0 | 29.3±1.2 | 18.3±1.5 | 2.0±0.0 |
| same size | 132.0±0.0 | 132.0±0.0 | 18±0 | 157.3±5.5 | 114.7±7.1 | 9.3±2.3 |
| everything | 396.7±0.6 | 317.3±0.6 | 53±0 | 441.0±10.0 | 292.7±11.0 | 31.3±2.5 |

The overall level of accurate classification predictions increases from 51.7% to 57.5% from the protocol 3 to Cartesian ddG Rosetta methods. We also note that over all charged residues the category predictions accuracy was 47.9% for protocol 3 and increases to over 56.1% with cartesian ddG. The Cartesian ddG algorithm is more broadly useful across any type of protein mutation, while previous methods had uneven applicability.

## Discussion

Here we describe a number of issues in previous benchmark sets used to assess the quality of protein stability prediction software. In particular we have found a lack of adequate experimental data being included for mutations involving charged residues.

Using these updated benchmarks we show that protein stability prediction tools in Rosetta vary widely across different types of mutation classes. In addition, given that this problem is pervasive throughout the field, it is likely that the reported accuracy of many methods for stability prediction may not reflect the diversity of possible mutation types. We encourage other developers to assess their tools on our benchmark set or on one which has appropriately accounted for the biases that exist within the databases (Supplemental Table 1).

Last we have refactored the Cartesian ddG protocol code and introduced new backbone sampling around mutations involving proline. These use Cartesian degrees of freedom and sampling instead of the Rosetta standard torsion space sampling (Rohl et al.). By tuning these algorithms with the new benchmark set, and focusing on improvements in previously underrepresented categories of mutations (e.g. uncharged to charged), we are able to achieve an algorithm with improved correlation to experimental values and drastically improved ability to correctly classify (stabilizing/destabilizing/neutral) a mutation. These new algorithms show the importance of diverse datasets in algorithm training, and the possibility for cross-fertilization between structure prediction methods and computational protein engineering methods.

## Methods

### Benchmark Set Pruning

To create our benchmark set, we began by making a copy of the curated ProTherm database(Ó Conchúir et al.) and began removing entries that were unsuitable. Because we wished to train a point mutation algorithm without the complexities of multiple mutation interactions, we excluded any entry which did not represent a single mutation. Because the trained algorithm is intended to represent ΔΔG of monomer folding and not binding interactions, we also removed entries on the interface of a protein-protein complex, or interacting with a non-water ligand. Interactions were defined as any atom in the mutated residue within 5 Å of an atom not on the same chain. To prevent overtraining, we wished to remove duplicate mutations. To identify duplicates, we performed an all to all sequence alignment to find parent backbones with ≥60% sequence identity. Within these clusters of sequences, any entries in which the same native residue is mutated to the same target were treated as identical. When multiple experimental ΔΔG values were available for an identical mutation we chose the value taken at closer to neutral pH.

### Benchmark category population

We identified 20 categories of mutation type by combinations from 9 residue type classifications (Table 3). We then populated each category with up to 50 entries. Mutually exclusive categories, i.e. negative to hydrophobic, were populated by picking a random entry from the curated databank and adding it to the benchmark if its respective category contained fewer than 50 entries. This was repeated 50,000 times after which we went through the entire databank from beginning to end populating any categories which were not already satisfied. Broad and non-exclusive categories, such as small to large, were sufficiently populated by the experiments selected from the exclusive group. A few categories involving charged residues (positive to negative, negative to positive, non-charged polar to positive, hydrophobic to negative, hydrophobic to positive, and like to like charge) did not have enough data to hit 50 entries so every available unique experiment was added.

**Table 3:** Residue category assignments and category combinations. On the left, we list the 9 residue groupings considered in this benchmark and annotate which residues go in each class. At center and right, we list the 20 mutational types considered by combining these classes.

| Type and 1 letter codes | Combination categories |  |
| --- | --- | --- |
| small GAVSTC | positive to negative | non-charged polar to hydrophobic |
| large FYWKRHQE | negative to positive | hydrophobic to non-charged polar |
| negative DE | positive to non-charged polar | non-charged polar to non-charged polar |
| positive RK | negative to non-charged polar | hydrophobic to hydrophobic |
| polar YTSHKREDQN | non-charged polar to positive | like to like charge |
| non-charged polar YTSNQH | non-charged polar to negative | involves proline |
| hydrophobic FILVAGMW | negative to hydrophobic | involves cysteine' |
| cysteine C | hydrophobic to negative | small to large |
| proline P | positive to hydrophobic | large to small |
|  | hydrophobic to positive | same size to same size |

### ΔΔG Prediction

To prepare models for ΔΔG calculations, structures were stripped to only the chain in which the mutation occurs. Rosetta local refinement (“Relax”) was then performed 20 times and the model with the lowest Rosetta energy was selected as input. ΔΔG predictions were then performed using protocol 3 described in Kellogg et al. 2010 or the modified version of the Cartesian ddG application described originally in Park et al. 2016 and elaborated upon here.

Modifications to the Cartesian ddG app include a change to the number of mutant models generated from a fixed number, 3, to a variable number based on the following convergence criterion: the lowest energy 2 structures must converge to within 1 Rosetta Energy Unit, or take the best of 5 models, whichever comes first. In order to address changes in the backbone resulting from mutations to and from proline we added additional fragment based sampling around mutations involving proline. By default 30 fragments of 5 residues in length are sampled and the best scoring structure is carried forward for analysis. This uses the Cartesian Sampler system described in Wang et al. (Wang et al.).

